# Generalized Chargaff symmetry in codon usage across the tree of life

**DOI:** 10.64898/2026.06.22.733775

**Authors:** Virginia Iannibelli, Isabella Caranzano, Giovanni Birolo, Cesare Rollo, Francesco Codicè, Guido Barducci, Cristian Taccioli, Luca Pagani, Piero Fariselli

**Affiliations:** AI and Computational Biomedicine Unit, Department of Biomedical Sciences, University of Torino, Torino, Italy; Department of Animal Medicine, Health and Production, University of Padova, Padova, Italy; Department of Biology, University of Padova, Padova, Italy

**Keywords:** codon usage, Chargaff symmetry, reverse complement, symmetry breaking, effective energy

## Abstract

Codon usage bias is a central record of mutation, selection, drift, and translational constraints, but it is usually treated separately from generalized Chargaff symmetry, the tendency for words and their reverse complements to occur at similar frequencies in long DNA sequences. Here we ask whether codon usage contains a measurable reverse-complement component, and whether departures from that component can be quantified. We first avoided imposing reverse-complement symmetry. Instead, we exhaustively evaluated all nontrivial *factorized codon involutions*. Across taxa, the reverse-complement transformation was the *optimum*, giving the highest median correlation between codon frequencies and transformed codon frequencies. Random and amino-acid-preserving reference models showed that the signal is not a generic property of codon profiles and is only partly explained by protein composition. Additional controls preserving amino-acid composition and matching GC3 in expectation showed that the observed reverse-complement correlation remains higher than expected from these constraints alone, and genus-level aggregation confirmed that the optimum is not driven by overrepresented genera. The symmetry breaks in a biologically ordered manner: the third, most degenerate codon position remains closest to the reverse-complement baseline, whereas the first departs most strongly and the second is intermediate and lineage dependent. Taxonomic comparisons reveal broad and fine-scale heterogeneity in codon-level symmetry preservation. Together, these results show that codon usage combines reverse-complement preservation with position-, lineage-, and function-dependent departures from the GCT-associated compositional baseline.

## Introduction

Coding sequences are embedded in DNA, whose double-stranded structure is associated with reverse-complement compositional symmetries. The message is familiar: coding sequences, regulatory elements, structural RNAs, and chromosomal features specify biological function. The symmetry is equally fundamental but less often placed at the center of genome analysis: double-stranded DNA couples every word to its reverse complement. If generalized Chargaff symmetry provides a compositional baseline, then coding information should appear not only as enrichment of particular motifs or codons, but also as structured departures from that baseline. This paper tests that idea at the scale of codon usage. Earlier work on sense–antisense coding and complementary oligonucleotide frequencies had already noted similarities between codon frequencies on coding and antisense strands, and discussed these patterns in relation to complementary-word parity and genomic sequence organization [1–3]. Building on this intuition, we ask a deliberately falsifiable question: without assuming reverse complement, does it emerge as the optimal codon-level transformation within the biologically interpretable class of factorized codon involutions?

Codon usage bias is a useful setting for this test because it connects sequence composition with evolutionary and translational constraints. Synonymous codons are not used equally, and their frequencies reflect mutation pressure, GC content, translational selection, tRNA abundance, gene expression, amino acid composition, and phylogenetic history [4–10]. These mechanisms are well established. What remains less clear is whether codon usage also retains a generalized Chargaff signature at the level of codon frequencies. Such a signal would not replace mutation, selection, drift, or biased fixation; it would provide a compositional reference against which their effects can be measured.

The relevant baseline comes from Chargaff’s second parity rule and its extension to longer words. On a single strand of long double-stranded DNA, complementary bases tend to occur at similar frequencies, A ≈T and C ≈G. Generalized Chargaff’s Theory (GCT) extends this regularity to *k*-mers: a word *w* and its reverse complement S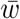 tend to have similar frequencies in long genomes [11, 12]. In the maximum-entropy formulation of [11], double-strandedness acts as a symmetry constraint: because every read of one strand is paired with a read of the complementary strand in the opposite direction, the effective sequence energy satisfies 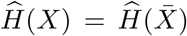, and the resulting equilibrium distribution gives 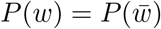 as the symmetric compositional baseline [11]. This effective energy is not a microscopic binding free energy. It is a coarse-grained statistical potential summarizing double-stranded constraints together with context-dependent mutation and repair, drift, selection, recombination-associated fixation biases, and other evolutionary processes over long timescales [12–14].

Those physical ideas were developed previously. The new question here is sharper and deliberately falsifiable: if codon usage is searched without assuming Chargaff symmetry, does the reverse complement emerge as the best codon-level symmetry? To answer this, we treated codon symmetry as a complete symbolic-search problem. We enumerated all nontrivial factorized codon involutions, defined as self-inverse codon transformations that decompose into a single self-inverse nucleotide map applied uniformly to the three codon positions, together with a self-inverse permutation of codon positions, and we asked which transformation maximizes the concordance between observed codon frequencies and transformed codon frequencies across taxa. This design makes the central result interpretable: reverse complement can win only if codon usage preserves the generalized Chargaff symmetry better than any alternative factorized base-and-position rule in the searched class.

Importantly, this test is not a direct restatement of generalized Chargaff parity for genomic 3-mers. Codon usage is a frame-specific projection of coding sequence, constrained by the genetic code, amino-acid demand, synonymous choice, and translational selection. Generalized Chargaff symmetry for whole-genome word frequencies, therefore does not require the 64 coding triplets, counted in their reading frame, to be optimally symmetric under reverse complement. The exhaustive search asks whether the reverse-complement symmetry associated with GCT remains detectable after this projection onto codon usage.

We then asked where this symmetry is preserved or lost. We compared observed codon profiles with reference models, decomposed the signal by codon position, examined taxonomic variation, and calibrated how small codon-pair effective-energy asymmetries can reduce reverse-complement concordance. The emerging picture is structured reverse-complement preservation and departure. The reverse-complement transformation provides a measurable compositional reference; the genetic code, lineage-specific evolution, and codon-level constraints can move codon usage away from it. Thus, codon usage shows measurable preservation of generalized Chargaff symmetry alongside departures associated with coding or evolutionary constraints.

## Results

### Reverse complement is selected by codon usage, not assumed

**Figure 1:**
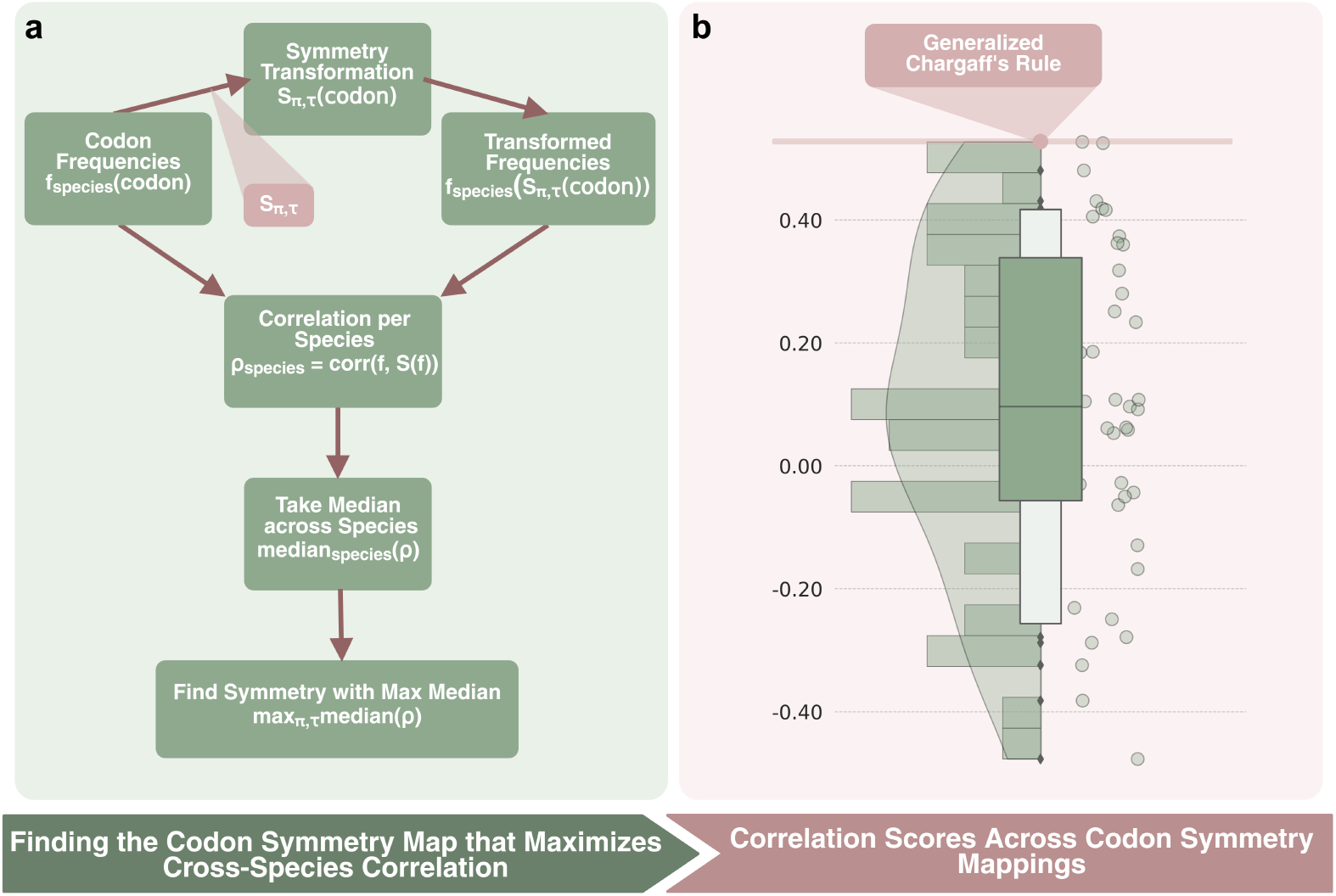
Exhaustive search over factorized codon involutions. We evaluated transformations *S*_*τ,π*_ defined by a self-inverse nucleotide map *τ* and a self-inverse codon-position permutation *π*. For each species, we computed the correlation between codon frequencies and transformed codon frequencies, *ρ*_species_ = corr(*f, S*(*f*)), and ranked transformations by the median correlation across species. The reverse-complement transformation maximized this criterion.

The title claim rests first on a simple test: reverse complement should not be inserted into the analysis if it can be selected from data. We therefore searched all factorized codon involutions, a biologically interpretable subclass of codon-level involutions. Each transformation *S* satisfied *S*(*S*(*c*)) = *c* for every codon *c*, preserving one-to-one correspondence and avoiding information loss. We restricted the rule space to transformations that act uniformly on codons by combining a nucleotide involution *τ* with a positional involution *π*:

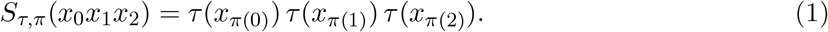

Here *τ* is a self-inverse nucleotide map applied uniformly to all three positions, and *π* is a self-inverse permutation of codon positions. This factorized construction gives 10 possible nucleotide involutions and 4 positional involutions, hence 40 transformations; after excluding the identity, 39 nontrivial candidate symmetries remained. The search is exhaustive within this factorized base-and-position class, not within the much larger set of arbitrary involutions on the 64-codon alphabet.

For each species *k*, we represented codon usage as a normalized vector *f*_*k*_ over the 64 codons and computed

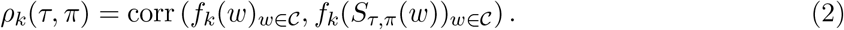

Appendix B gives a mathematical interpretation of this statistic: for any codon-level involution, maximizing the Pearson correlation is equivalent to minimizing the normalized component of codon usage that is odd under that involution. We then ranked each transformation by the median value of *ρ*_*k*_(*τ, π*) across species. This exhaustive search within the factorized base-and-position rule space identified a single standout optimum: Watson–Crick complementation, A ↔ T and C ↔ G, combined with reversal of the three codon positions, *π* = (2, 1, 0). This is precisely the codon reverse complement.

The reverse-complement transformation achieved the highest median Pearson correlation across species, *ρ* = 0.53, substantially above the distribution median across all candidate mappings (*ρ* = 0.10; first quartile = −0.06, third quartile = 0.34). The second-best transformation used the identity positional permutation and a nucleotide map that swapped only G and C, consistent with a strong contribution of GC composition. Nevertheless, it did not exceed reverse complement. Thus, when codon usage is viewed as a symbolic symmetry problem, the reverse complement is not inserted by theory; it is selected by the data.

This optimum is conservative with respect to trivial self-matching. Several alternative involutions contain fixed codons, so that 8 or 16 of the 64 codon-frequency entries are compared with themselves when computing *ρ*_*k*_(*τ, π*). Such fixed points can mechanically increase a Pearson correlation. By contrast, the reverse-complement transformation has no fixed codons and maps no sense codon to a synonymous codon, yet it remains the top-ranked transformation (Appendix B.4 in Table 1).

**Table 1:**
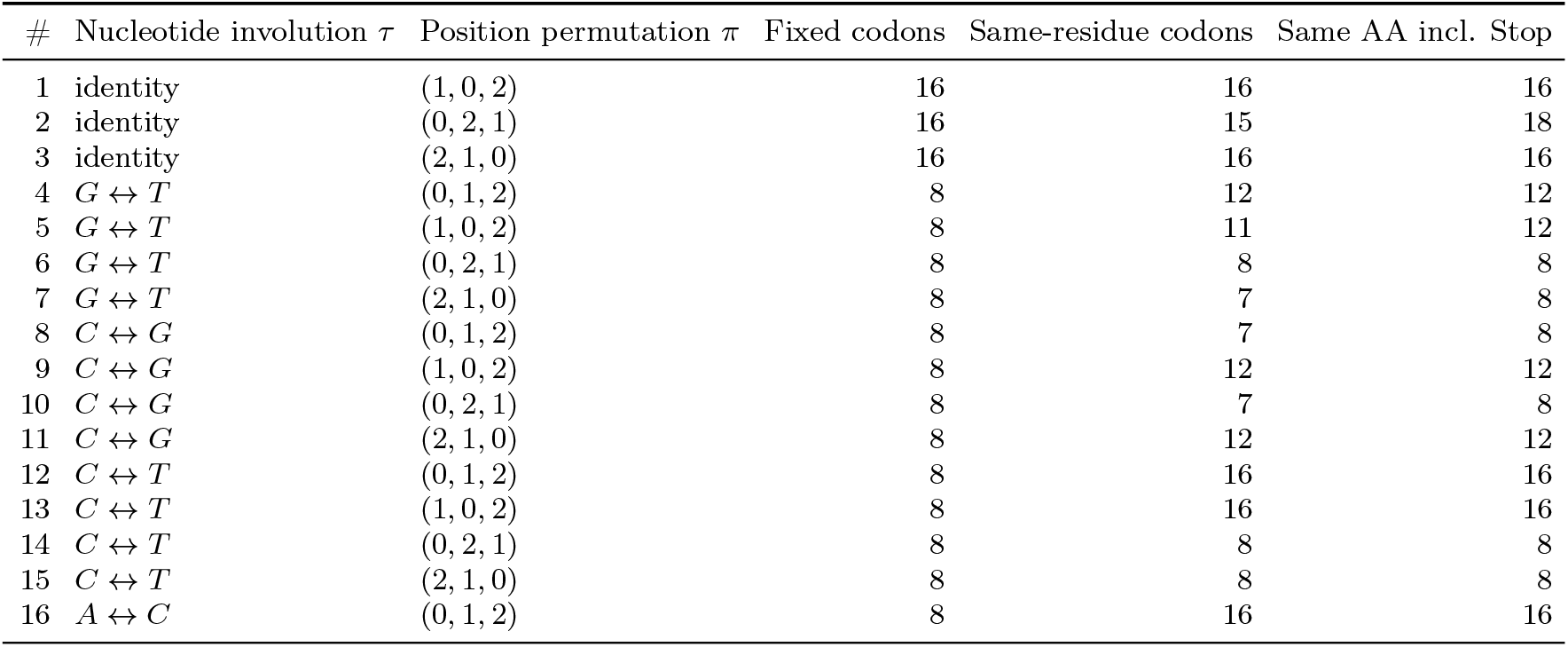
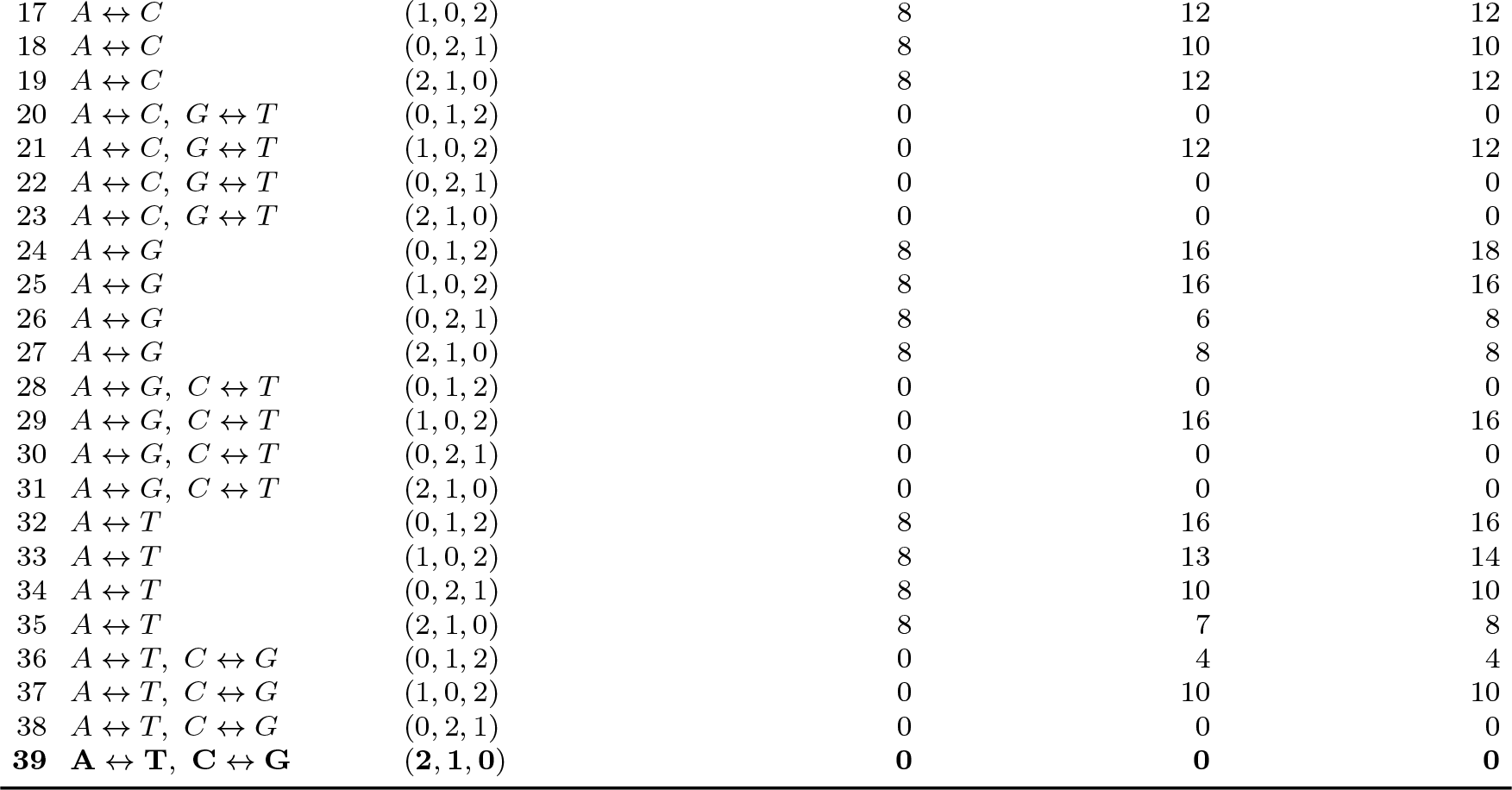
Fixed codons and amino-acid preservation for the 39 non-identity factorized codon involutions. Fixed codons are codons *c* satisfying *S*(*c*) = *c*. Same-residue codons count sense codons for which *a*(*S*(*c*)) = *a*(*c*), excluding stop codons as residues. The last column treats stop as an additional amino-acid class. The reverse-complement transformation is row 39.

| # | Nucleotide involution $\tau$ | Position permutation $\pi$ | Fixed codons | Same-residue codons | Same AA incl. Stop |
| --- | --- | --- | --- | --- | --- |
| 1 | identity | (1, 0, 2) | 16 | 16 | 16 |
| 2 | identity | (0, 2, 1) | 16 | 15 | 18 |
| 3 | identity | (2, 1, 0) | 16 | 16 | 16 |
| 4 | $G \leftrightarrow T$ | (0, 1, 2) | 8 | 12 | 12 |
| 5 | $G \leftrightarrow T$ | (1, 0, 2) | 8 | 11 | 12 |
| 6 | $G \leftrightarrow T$ | (0, 2, 1) | 8 | 8 | 8 |
| 7 | $G \leftrightarrow T$ | (2, 1, 0) | 8 | 7 | 8 |
| 8 | $C \leftrightarrow G$ | (0, 1, 2) | 8 | 7 | 8 |
| 9 | $C \leftrightarrow G$ | (1, 0, 2) | 8 | 12 | 12 |
| 10 | $C \leftrightarrow G$ | (0, 2, 1) | 8 | 7 | 8 |
| 11 | $C \leftrightarrow G$ | (2, 1, 0) | 8 | 12 | 12 |
| 12 | $C \leftrightarrow T$ | (0, 1, 2) | 8 | 16 | 16 |
| 13 | $C \leftrightarrow T$ | (1, 0, 2) | 8 | 16 | 16 |
| 14 | $C \leftrightarrow T$ | (0, 2, 1) | 8 | 8 | 8 |
| 15 | $C \leftrightarrow T$ | (2, 1, 0) | 8 | 8 | 8 |
| 16 | $A \leftrightarrow C$ | (0, 1, 2) | 8 | 16 | 16 |
| 17 | $A \leftrightarrow C$ | (1, 0, 2) | 8 | 12 | 12 |
| 18 | $A \leftrightarrow C$ | (0, 2, 1) | 8 | 10 | 10 |
| 19 | $A \leftrightarrow C$ | (2, 1, 0) | 8 | 12 | 12 |
| 20 | $A \leftrightarrow C, G \leftrightarrow T$ | (0, 1, 2) | 0 | 0 | 0 |
| 21 | $A \leftrightarrow C, G \leftrightarrow T$ | (1, 0, 2) | 0 | 12 | 12 |
| 22 | $A \leftrightarrow C, G \leftrightarrow T$ | (0, 2, 1) | 0 | 0 | 0 |
| 23 | $A \leftrightarrow C, G \leftrightarrow T$ | (2, 1, 0) | 0 | 0 | 0 |
| 24 | $A \leftrightarrow G$ | (0, 1, 2) | 8 | 16 | 18 |
| 25 | $A \leftrightarrow G$ | (1, 0, 2) | 8 | 16 | 16 |
| 26 | $A \leftrightarrow G$ | (0, 2, 1) | 8 | 6 | 8 |
| 27 | $A \leftrightarrow G$ | (2, 1, 0) | 8 | 8 | 8 |
| 28 | $A \leftrightarrow G, C \leftrightarrow T$ | (0, 1, 2) | 0 | 0 | 0 |
| 29 | $A \leftrightarrow G, C \leftrightarrow T$ | (1, 0, 2) | 0 | 16 | 16 |
| 30 | $A \leftrightarrow G, C \leftrightarrow T$ | (0, 2, 1) | 0 | 0 | 0 |
| 31 | $A \leftrightarrow G, C \leftrightarrow T$ | (2, 1, 0) | 0 | 0 | 0 |
| 32 | $A \leftrightarrow T$ | (0, 1, 2) | 8 | 16 | 16 |
| 33 | $A \leftrightarrow T$ | (1, 0, 2) | 8 | 13 | 14 |
| 34 | $A \leftrightarrow T$ | (0, 2, 1) | 8 | 10 | 10 |
| 35 | $A \leftrightarrow T$ | (2, 1, 0) | 8 | 7 | 8 |
| 36 | $A \leftrightarrow T, C \leftrightarrow G$ | (0, 1, 2) | 0 | 4 | 4 |
| 37 | $A \leftrightarrow T, C \leftrightarrow G$ | (1, 0, 2) | 0 | 10 | 10 |
| 38 | $A \leftrightarrow T, C \leftrightarrow G$ | (0, 2, 1) | 0 | 0 | 0 |
| **39** | **$A \leftrightarrow T, C \leftrightarrow G$** | **(2, 1, 0)** | **0** | **0** | **0** |

To reduce the effect of uneven taxonomic sampling, we repeated the factorized-codon-involution ranking after collapsing species-level correlations at the genus level. Across 4,445 genera, the reverse-complement transformation remained the top-ranked map (median genus-level *ρ* = 0.5157), above the best non-reverse-complement transformation (median *ρ* = 0.4814; margin = 0.0343). In 1,000 bootstrap replicates in which one species was sampled at random from each genus, the reverse-complement transformation ranked first in 1,000/1,000 replicates, with median margin 0.0382 over the best non-reverse-complement map.

### Reference models show the signal is not a random codon-profile artifact

A symmetry baseline is useful only if it is not a trivial consequence of arbitrary codon profiles. We therefore asked whether the observed reverse-complement signal could be explained by simple properties of codon usage (Figure 2). For each species, we computed the codon-level reverse-complement correlation 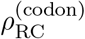, defined as the Pearson correlation between the 64 codon frequencies and the corresponding reverse-complement codon frequencies. We compared the observed values with three reference models. In the random model, codons were assigned independent random values and renormalized. In the amino-acid-preserving model, codon frequencies were shuffled within synonymous codon families, preserving amino acid composition but disrupting synonymous codon preference. In the normal reference model, values were sampled from a Gaussian distribution calibrated to the random model.

**Figure 2:**
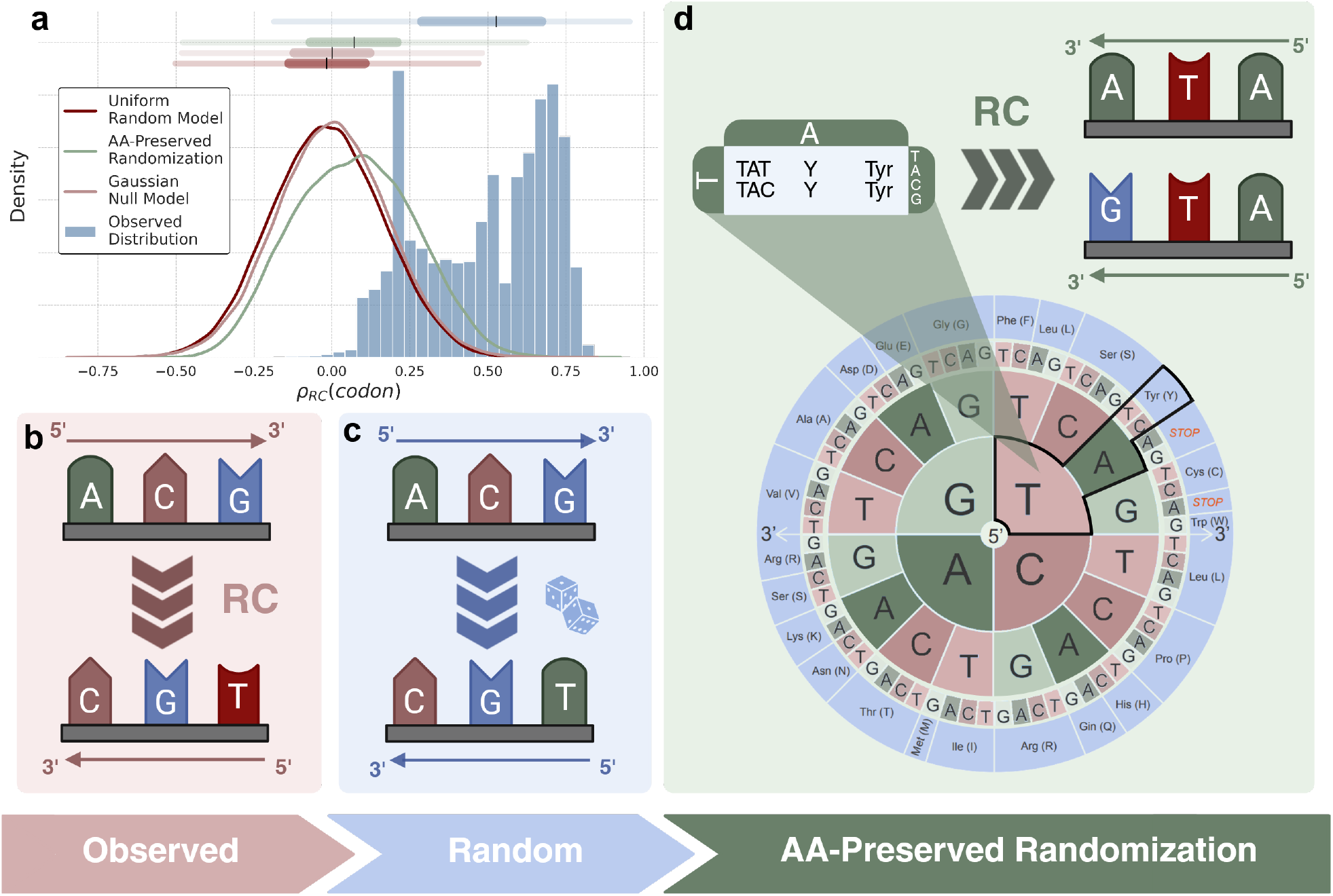
Codon-level reverse-complement correlations in observed and reference profiles. The observed distribution of 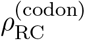 is compared with random codon profiles, amino-acid-preserving shuffles of codon usage within synonymous groups, and a normal reference distribution based on the variability of the random model (panel **a**). **Observed** represents the distribution of correlations from the experimental dataset (panel **b**). **Random** indicates the correlation computed after assigning uniform random values to the codon usage frequencies, and **Normal** is the distribution sampled from a Gaussian with zero mean and a standard deviation equal to that observed in the Random model (panel **c**). **AA-Preserved Randomization** corresponds to the distribution obtained by randomly shuffling the codon usage frequencies within each group of synonymous codons (thus preserving overall amino acid composition, see panel **d**).

**Figure 3:**
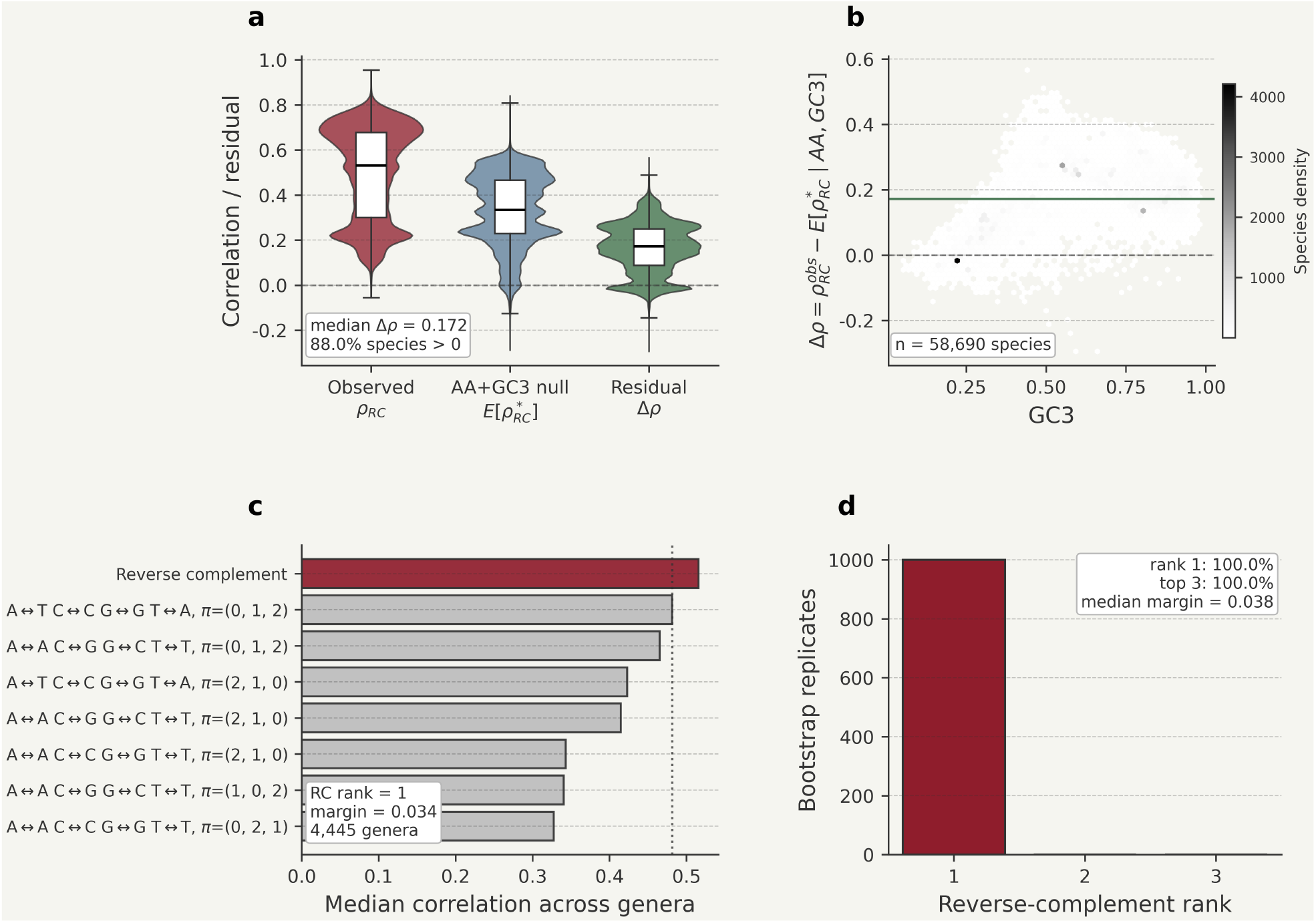
AA+GC3 and genus-level controls for codon reverse-complement symmetry. **a**, Observed codon-level reverse-complement correlations, 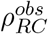, are compared with an amino-acid- and GC3-controlled synonymous-codon null model. For each species, the null model preserves amino-acid composition and matches the observed third-position GC content (GC3) in expectation, while otherwise randomizing synonymous codon choice. The residual signal is defined as 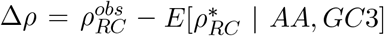. Across 58,690 species with at least 100,000 codons, the observed correlations remained higher than the AA+GC3 expectation (median 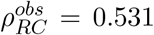, median 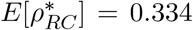, median Δ*ρ* = 0.172); 88.0% of species had Δ*ρ >* 0. **b**, Residual reverse-complement correlation as a function of GC3. Positive residuals across a broad GC3 range indicate that amino-acid composition and third-position GC content explain part, but not all, of the codon-level reverse-complement signal. **c**, Genus-level robustness of the exhaustive factorized-codon-involution ranking. Species-level correlations were collapsed by genus using the median, and transformations were ranked by their median correlation across 4,445 genera. The reverse-complement transformation remained the top-ranked factorized codon involution, with a median genus-level correlation of 0.516 and a margin of 0.0343 over the best non-reverse-complement transformation. **d,** One-species-per-genus bootstrap analysis. In each of 1,000 bootstrap replicates, one species was sampled from each genus and the 39 factorized codon involutions were re-ranked. The reverse-complement transformation ranked first in 1,000/1,000 replicates, showing that the global optimum is not driven by overrepresented genera.

The random and normal reference distributions were centered near zero, showing that arbitrary codon profiles do not generate systematic reverse-complement correlations. Amino-acid-preserving shuffles produced a modest positive shift, indicating that protein-level composition and degeneracy can contribute to reverse-complement balance. However, the observed distribution was displaced beyond these simple baselines (Figure 2). This result is consistent with at least two additive contributions to reverse-complement balance: one arising from protein composition and synonymous degeneracy, and a further component in observed codon usage that persists after controlling for amino-acid content and is consistent with the compositional background expected under generalized Chargaff symmetry.

Because GC3 is a relevant determinant of codon usage, we next asked whether the reverse-complement signal could be explained by amino-acid composition and third-position GC content alone. For each species with at least 100,000 codons (*n* = 58,690), we constructed an amino-acid- and GC3-controlled synonymous-codon null model that preserves amino-acid composition and matches the observed GC3 in expectation. The observed reverse-complement correlations remained substantially higher than this null expectation (median observed 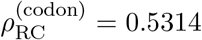; median AA+GC3 null 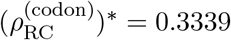; median Δ*ρ* = 0.1717), with 88.0% of species showing Δ*ρ >* 0 (Wilcoxon signed-rank test, *p* below numerical precision). After collapsing species-level results by genus, the excess remained positive in 94.0% of 4,445 genera, with median genus-level Δ*ρ* = 0.2023. Thus, amino-acid composition and GC3 account for part, but not all, of the codon-level reverse-complement signal.

These GC3-controlled and genus-level robustness analyses are summarized in Figure.

### Functional symmetry breaking is ordered within the codon

If reverse-complement balance provides a baseline, then coding constraints should affect the three codon positions differently. The codon-level correlation compresses positions with different biological roles. We therefore decomposed the signal into adjacent two-position blocks and single codon positions (Fig. 4). For the two-position analysis, codon frequencies were summed over the remaining nucleotide to obtain frequencies for positions 1–2 and 2–3. A pair (*a, b*) was compared with its reverse complement 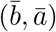. For the single-position analysis, we summed over the other two codon positions to obtain the frequency of each nucleotide at positions 1, 2, and 3, and compared each distribution with its complementary-base distribution.

**Figure 4:**
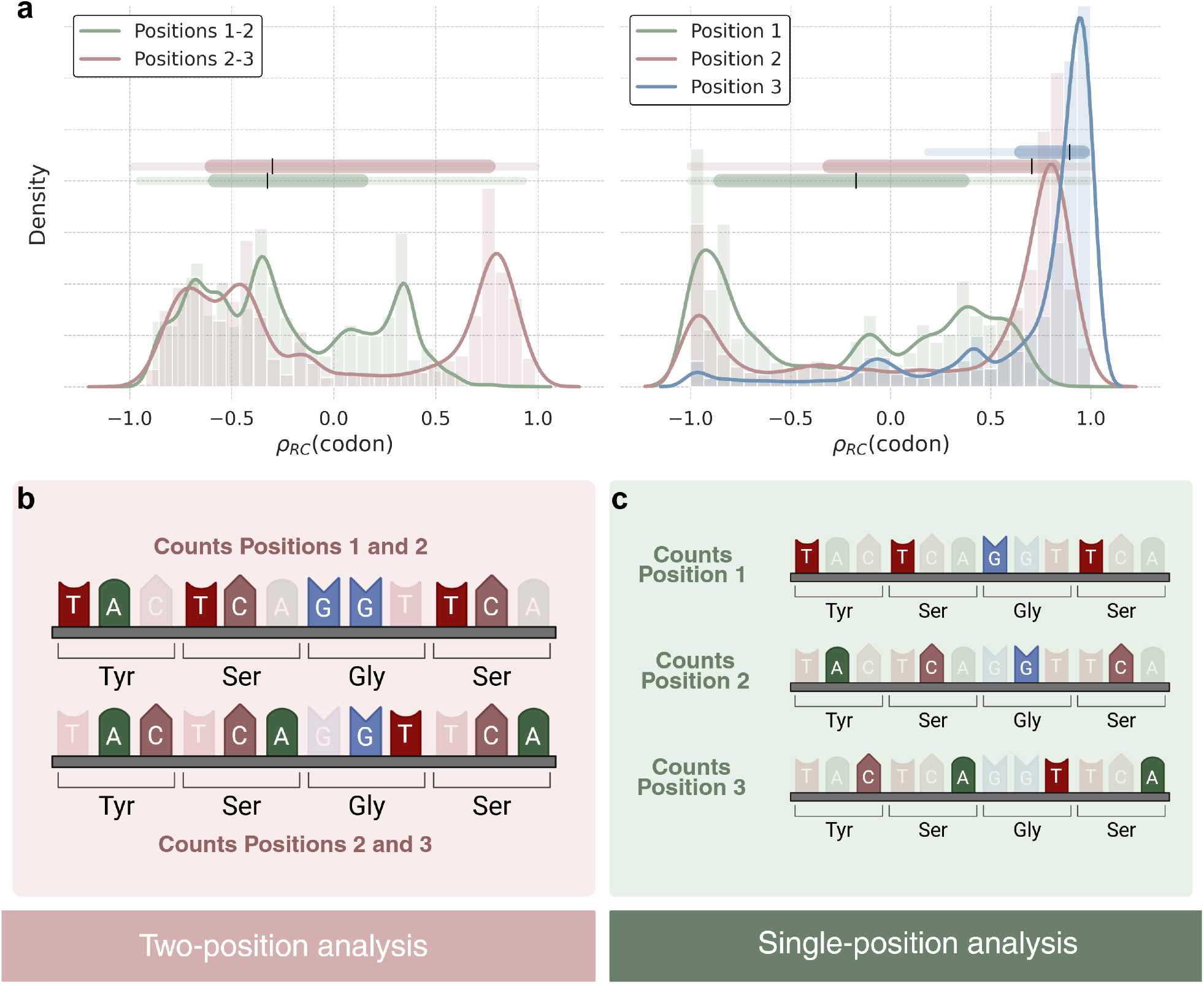
Reverse-complement symmetry at the subcodon level. **a** Distributions across genomes of correlation values in the two-position and single-position analyses. **b** Two-position analysis: codon frequencies were aggregated into positions 1–2 and 2–3 by summing over the remaining base, and the two blocks were compared with their reverse complements. **c** Single-position analysis: nucleotide frequencies were aggregated at codon positions 1, 2, and 3 and compared with complementary bases. Only species above the codon-count threshold were included.

The two-position analysis revealed opposite tendencies for the two adjacent windows. Frequencies aggregated over positions 1–2 were shifted toward negative correlations, whereas those aggregated over positions 2–3 were shifted toward positive correlations. The single-position analysis resolved this contrast. The first codon position showed the strongest and most consistent departure from reverse-complement symmetry. The second position was intermediate, with a broad distribution and substantial cross-species variability. The third position, corresponding to the wobble site, showed a narrow distribution concentrated near *ρ*_RC_ = 1, indicating strong adherence to the reverse-complement baseline. Thus, symmetry breaking is not random within codons: it is largest at position 1, intermediate and lineage-dependent at position 2, and smallest at the wobble position where degeneracy allows greater compositional freedom. The strong departure at position 1 likely reflects the combination of translational selection and context-dependent mutational biases acting on the first base, while the breadth of the position-2 distribution may reflect the fact that changes at this position are the most biochemically conservative on average, leaving greater lineage-specific latitude.

### Lineages differ in preservation of generalized Chargaff symmetry

Functional symmetry breaking should also depend on evolutionary history. Consistent with this view, the distribution of 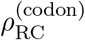 changed markedly across taxonomic depth (Fig. 5). At the broadest level, Archaea and Bacteria showed the highest median values, 0.545 and 0.532, respectively. Eukaryota had a lower median value of 0.459, and Viruses were lower still, with a median of 0.382. These top-level groups differed strongly overall (Kruskal–Wallis *H* = 1640.57, *p <* 10^*−*300^), and all pairwise comparisons were significant by Mann–Whitney tests (*p <* 0.001).

**Figure 5:**
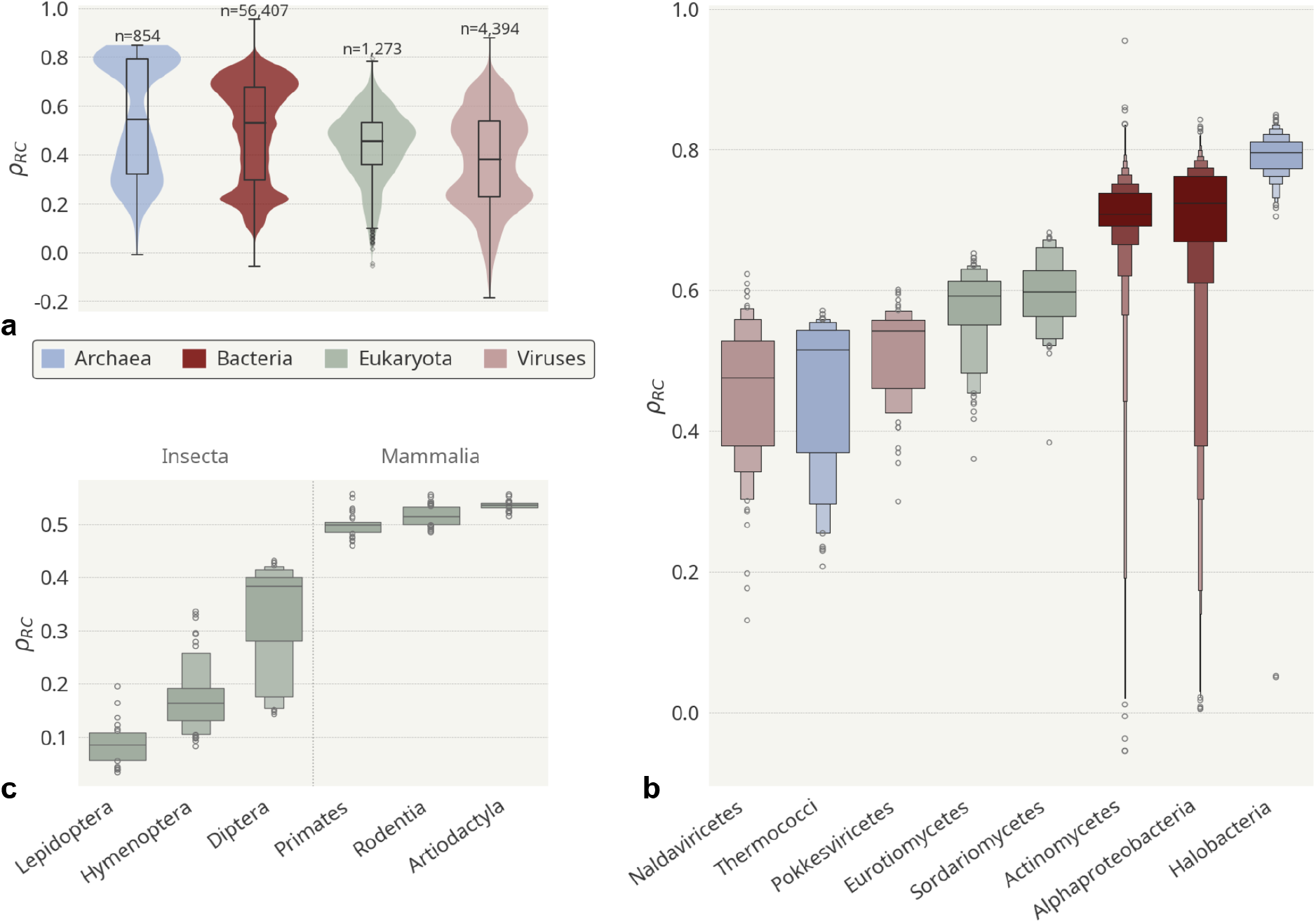
Distribution of 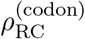 across taxonomic levels. **a** Violin and box plots for Archaea, Bacteria, Eukaryota, and Viruses. Group differences were assessed with a Kruskal–Wallis test (*H* = 1640.57, *p <* 10^*−*300^), followed by pairwise Mann–Whitney tests, all significant at *p <* 0.001. **b** Class-level distributions for selected representative classes; group differences were significant by Kruskal–Wallis test (*H* = 1808.05, *p <* 10^*−*300^). **c** Order-level comparison of selected insect and mammalian orders. Lepidoptera differed significantly from each mammalian order (*p <* 0.001), and pooled insect and mammalian groups differed in dispersion by Levene’s test (*p* = 3.22 × 10^*−*20^; *n* = 23–64 per order).

At class level, the broad pattern resolved into more pronounced lineage-specific differences. Halobacteria showed the highest median value among the selected classes (0.796), followed by Alphaproteobacteria (0.724) and Actinomycetes (0.708), whereas viral and fungal classes occupied lower parts of the distribution, including Naldaviricetes (0.475), Eurotiomycetes (0.592), and Sordariomycetes (0.598). At order level, insect and mammalian groups contrasted sharply. Lepidoptera and Hymenoptera had low median values, 0.085 and 0.163, respectively, whereas mammalian orders clustered at higher values: Primates at 0.498, Rodentia at 0.513, and Artiodactyla at 0.536. The mammalian orders were comparatively tight, whereas insect orders spanned a wider range. These patterns indicate that codon-level reverse-complement symmetry is neither uniformly conserved nor randomly distributed. Instead, it is modulated by lineage-specific combinations of base composition, mutation and repair biases, recombination-associated fixation biases, genome architecture, and coding constraints.

Together, these comparisons show that the taxonomic differences are superimposed on a broader reverse-complement signal. Because random and normal codon profiles are centered near zero, the positive values observed across major groups indicate that codon usage is generally displaced toward the reverse-complement balance expected under GCT rather than being randomly organized. The main taxonomic effect is therefore not the presence or absence of this baseline, but the degree to which it is preserved or broken. Archaea and Bacteria remain comparatively close to the reverse-complement baseline expected under generalized Chargaff symmetry, whereas viruses, eukaryotes, and especially some insect orders show stronger lineage-specific departures. Thus, taxonomic structure is itself part of the symmetry-breaking pattern, consistent with different evolutionary regimes modulating a shared generalized Chargaff baseline.

A phylum-resolved view of reverse-complement correlations across all domains is provided in Fig. S1 (SI Appendix), and an interactive version for searching and ranking phyla is available at https://compbiomed-unito.github.io/chargaff-symmetry-tradeoffs/.

### Small effective asymmetries are sufficient to break codon symmetry

Using this codon-level calibration, assigning an effective asymmetry of approximately 0.3 *k*_*B*_*T* to one member of each codon–reverse-complement pair is sufficient to reduce a near-perfect symmetry correlation to a value close to that observed for human codon usage (*ρ* ≈ 0.50 in Fig. 6).

**Figure 6:**
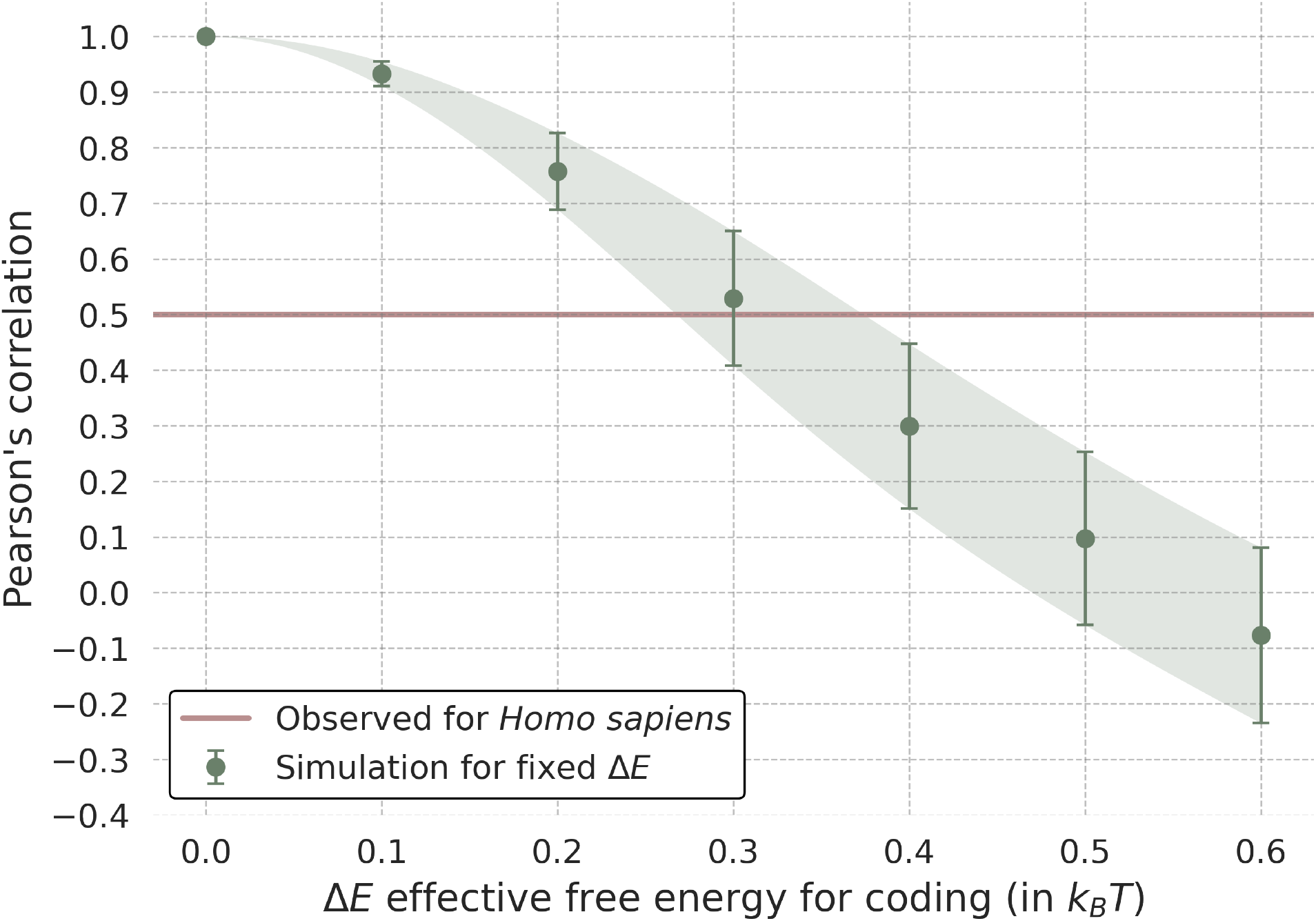
Effective-energy calibration of reverse-complement correlation. Simulated codon frequencies were generated by assigning shared baseline energies to codon–reverse-complement pairs and adding an asymmetric cost Δ*E* to one member of each pair. Increasing Δ*E* reduces the expected reverse-complement Pearson correlation. The value 0.3 *k*_*B*_*T* is shown only as an order-of-magnitude reference to the coding-associated scale estimated from 7-mer effective energies in human chromosomes [12]; no direct conversion to a codon-level cost is assumed.

This scale is broadly consistent with previous estimates of coding-associated effective cost in human chromosomes derived from 7-mer analyses [12], which were also of order 0.3 *k*_*B*_*T*. Although the two analyses operate at different sequence scales (7-mer frequencies versus codon-level, *k* = 3, reverse-complement pairs), the numerical concordance suggests that both approaches may be sensitive to comparable effective asymmetries associated with coding sequence. In the present codon-pair calibration, an asymmetry of this magnitude is sufficient to reduce reverse-complement correlation to values close to those observed empirically, supporting the interpretation that modest effective biases, accumulated across codons, can appreciably break codon-level reverse-complement concordance.

## Discussion

This study makes codon usage readable as symmetry breaking relative to a generalized Chargaff baseline. The central finding is not a new derivation of Generalized Chargaff’s Theory and not a new effective-energy formalism; both were developed previously [11, 12]. The advance is that reverse complement emerges from an exhaustive search over factorized codon symmetries. Among all factorized codon involutions formed by a uniform nucleotide involution and a codon-position involution, the reverse complement maximizes codon-usage concordance across taxa. Thus the reverse-complement transformation associated with generalized Chargaff symmetry is not merely compatible with codon usage. It is the best-preserved codon-level symmetry in the complete factorized rule space considered here.

This changes the role of Chargaff symmetry in codon-usage analysis. Reverse complement is often treated as a known global property of DNA composition. Here it becomes a data-selected transformation at the scale of the genetic code. The test is deliberately falsifiable: if codon usage were organized primarily by another symbolic balance, another factorized base-and-position involution would have ranked higher. Instead, the reverse-complement transformation was the optimum, and its departures were biologically structured.

The taxonomic analysis adds an evolutionary dimension to the same principle. Prokaryotes have high median codon-level reverse-complement symmetry, viruses are lower, and finer-scale groups show substantial heterogeneity. These differences do not imply that some lineages obey physical constraints and others do not. Rather, they show that different evolutionary regimes superimpose different biases on the same broad compositional baseline. GC-biased gene conversion, context-dependent mutation, repair asymmetries, translational selection, compact genome architecture, overlapping coding requirements, and effective population size can all push codon usage away from reverse-complement balance in lineage-specific ways.

The low codon-level reverse-complement correlations observed in Lepidoptera and Hymenoptera are intriguing and may point to insect-specific departures from the generalized Chargaff compositional baseline. One speculative possibility is that ecological specialization contributes to this pattern. Lepidoptera, in particular, are a classical model of plant–insect coevolution [15], and host-associated adaptation could increase lineage-specific functional or translational constraints on coding sequences. However, the present analysis cannot distinguish this hypothesis from alternative explanations.

As a complementary analysis, Appendix A reports a genome-scale effective-energy landscape analysis of *Drosophila melanogaster*. This analysis asks whether regions enriched in statistically unexpected 7-mers relative to their chromosome background coincide with functional annotation; it is conceptually linked to the maximum-entropy formulation of generalized Chargaff symmetry, while remaining distinct from the codon-level transformation search. Windows with high average effective energy, i.e. enriched in chromosome-rare 7-mers, are strongly enriched in coding and other genic annotations, whereas low-scoring windows are predominantly intergenic or weakly annotated. This result is analogous to previous observations in human chromosomes but occurs in a compact invertebrate genome with different gene density and architecture [12]. This suggests that local effective-energy deviations may provide an additional marker of functional sequence composition across broadly different genomic contexts.

Several limitations remain. First, the effective energy used here is inferred from sequence frequencies, not from a direct physical measurement. More broadly, the reference models control for arbitrary codon profiles, amino-acid composition, and, in the AA+GC3 analysis, third-position GC content in expectation. Additional controls preserving dinucleotide composition, expression level, and codon adaptation would further refine the mechanism.

Despite these limitations, the conclusion is simple. Generalized Chargaff symmetry supplies a reverse-complement compositional reference. The genetic code, genome architecture, and lineage-specific evolutionary processes preserve or displace that reference in organized ways. Codon usage bias is therefore not only a record of mutation and selection, but also reflects how coding information is organized relative to reverse-complement symmetry.

## Materials and Methods

### Codon usage data

Codon usage data were obtained from the HIVE-CUTs database [16], which compiles codon counts from GenBank and RefSeq protein-coding sequences [17]. For each organism, absolute counts for all 64 codons were normalized by the total number of codons to obtain relative frequencies freq(*c*). We retained genomes using translation tables 0, 1, or 11 and containing at least 10,000 codons.

### Codon-level reverse-complement correlation

For each organism, the reverse complement of a codon *c* was obtained by reversing the codon and complementing each base, A↔T and C↔G. We defined

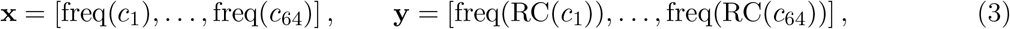

and computed *ρ*^(codon)^ = corr(**x, y**) using the Pearson correlation coefficient.

### Reference models

We used three reference models. In the random model, each codon was assigned a random value sampled uniformly from [0, 1], followed by normalization to sum to one. In the amino-acid-preserving model, codon frequencies were randomly permuted within each synonymous codon family, preserving amino acid composition while removing codon preference within each family. In the normal reference model, values were sampled from a Gaussian distribution with zero mean and a standard deviation matching the random model.

### AA+GC3-controlled reference model and genus-level robustness

To test whether the reverse-complement signal could be explained by amino-acid composition and third-position GC content, we constructed an AA+GC3 null model for each species. Let *N*_*a*_ be the observed number of codons assigned to amino acid *a*, let *N* = Σ_*a*_ *N*_*a*_, and let _*a*_ be the synonymous codon family for *a*. For codon *c*, define *I*_3_(*c*) = 1 if the third base is G or C and *I*_3_(*c*) = 0 otherwise. Conditional on amino acid *a*, the null probability of codon *c* ∈ C_*a*_ was

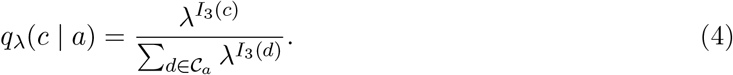

The parameter *λ* was fitted separately for each species so that the expected third-position GC content matched the observed value. The observed GC3 for species *k* was

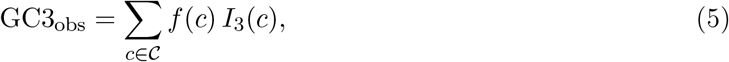

where C denotes the full set of 64 codons. The fitted value of *λ* was obtained by solving

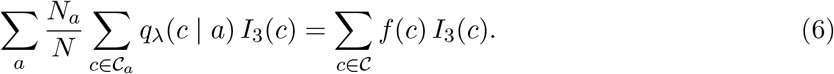

The two sides are numerically equivalent when amino-acid frequencies are derived from the same codon table, 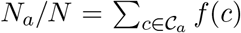, but writing the right-hand side explicitly fixes the definition of GC3_obs_ as a codon-frequency-weighted mean over all 64 codons.

The joint AA+GC3 null frequency over all codons was

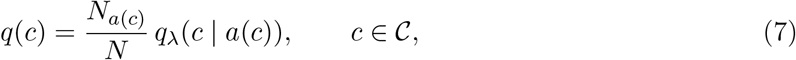

where *a*(*c*) ∈ {Phe, Leu, …, Stop} is the amino acid, or stop signal, encoded by codon *c*. Stop codons {TAA, TAG, TGA} were retained as a three-member synonymous family and treated identically to sense-codon families throughout. The fitted value of *λ* therefore also accounts for the third-position composition of stop codons.

Randomized profiles were generated by multinomial sampling within each synonymous family,

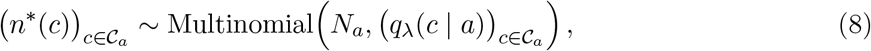

and the randomized null frequency profile was *f*^*∗*^(*c*) = *n*^*∗*^(*c*)*/N*. We summarized the controlled excess as

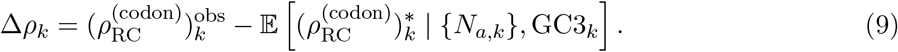

To reduce sampling bias from overrepresented genera, species-level results were also collapsed by genus using the median value within each genus. For the factorized-codon-involution ranking, we computed the genus-level statistic

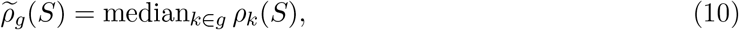

and ranked transformations by median 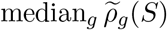. As a complementary robustness analysis, we performed 1,000 bootstrap replicates in which one species was selected at random from each genus and the full 39-transformation ranking was recomputed.

### Subcodon analyses

For the two-position analysis, codon frequencies were aggregated as

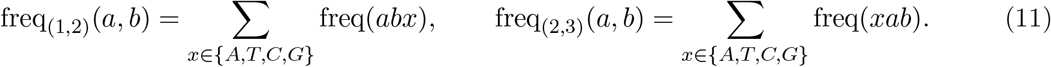

We compared freq_(1,2)_(*a, b*) with freq_(2,3)_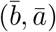 across the 16 possible pairs. For the single-position analysis, we computed

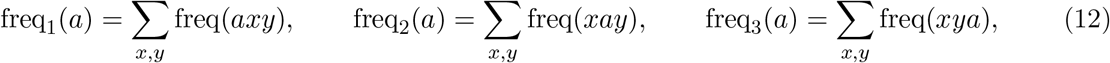

where *x, y* ∈ {*A, T, C, G*}, and compared each position-specific nucleotide distribution with the corresponding complementary-base distribution.

### Taxonomic annotation and statistics

For each organism, 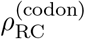 was stratified by broad taxonomic group and by lower-rank categories where available. HIVE-CUTs entries were merged with RefSeq and NCBI taxonomy metadata [17, 18]. Group differences were evaluated with Kruskal–Wallis tests, pairwise Mann–Whitney tests, and Levene’s test for dispersion where indicated. Exact test statistics are reported in the corresponding figure captions and Results sections.

### Optimization of factorized codon involutions

Because the full set of involutions on 64 elements exceeds 10^45^, we restricted the search to the factorized class of 39 transformations, each defined by a uniform base-exchange rule and a codon-position permutation and therefore directly interpretable in molecular terms. Let *C* = {*x*_0_*x*_1_*x*_2_ | *x*_*i*_ ∈ {*A, C, G, T* }} be the codon set. Each candidate transformation was defined as

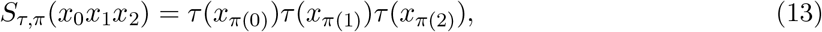

where *τ* is a nucleotide involution and *π* is a position involution. We evaluated all 39 non-identity transformations in this factorized class. This class is a restricted, biologically interpretable subset of all possible involutions on the 64-codon alphabet: each map is defined by a uniform base-exchange rule and a codon-position permutation, rather than by an arbitrary pairing of codons. For each transformation and species *k*, we computed *ρ*_*k*_(*τ, π*) as the Pearson correlation between the original codon frequency vector and the transformed frequency vector. The objective was

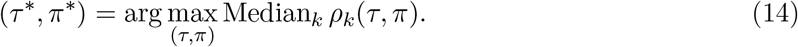

### Effective-energy calibration

For the codon-level calibration, codon–reverse-complement pairs were assigned shared baseline energies. An additional cost Δ*E* was added to one member of each pair with probability 1*/*2. Boltzmann-like frequencies were computed and normalized across the 64 codons, and the Pearson correlation between *P* (*c*) and *P* (RC(*c*)) was calculated as a function of Δ*E*. This analysis was used only to calibrate the scale of effective asymmetry required to reproduce observed codon-level correlations.

## A Genome-scale effective-energy landscape in *Drosophila melanogaster*

The codon-level analyses in the main text show that reverse complement is the best-preserved symbolic symmetry of codon usage across taxa, and that departures from this reference are organized by codon position and evolutionary lineage. We next asked whether the same effective-energy perspective provides a genome-scale readout of functional sequence organization. To this end, we analyzed the compact and well-annotated genome of *Drosophila melanogaster*, using chromosome-wide 7-mer frequencies to define a local effective-energy landscape.

This analysis extends the codon-scale results in two ways. First, it tests whether regions enriched in statistically unexpected words relative to their chromosome background preferentially coincide with functional annotations. Second, it connects the codon-level symmetry framework to the broader maximum-entropy formulation of generalized Chargaff symmetry, in which word frequencies define a coarse-grained sequence potential. Thus, the analysis is not used to rank codon transformations, but to ask whether effective-energy deviations from the chromosome-wide compositional background carry biological information at larger genomic scales.

### A.1 Effective-energy landscape marks functional sequence

**Figure A1:**
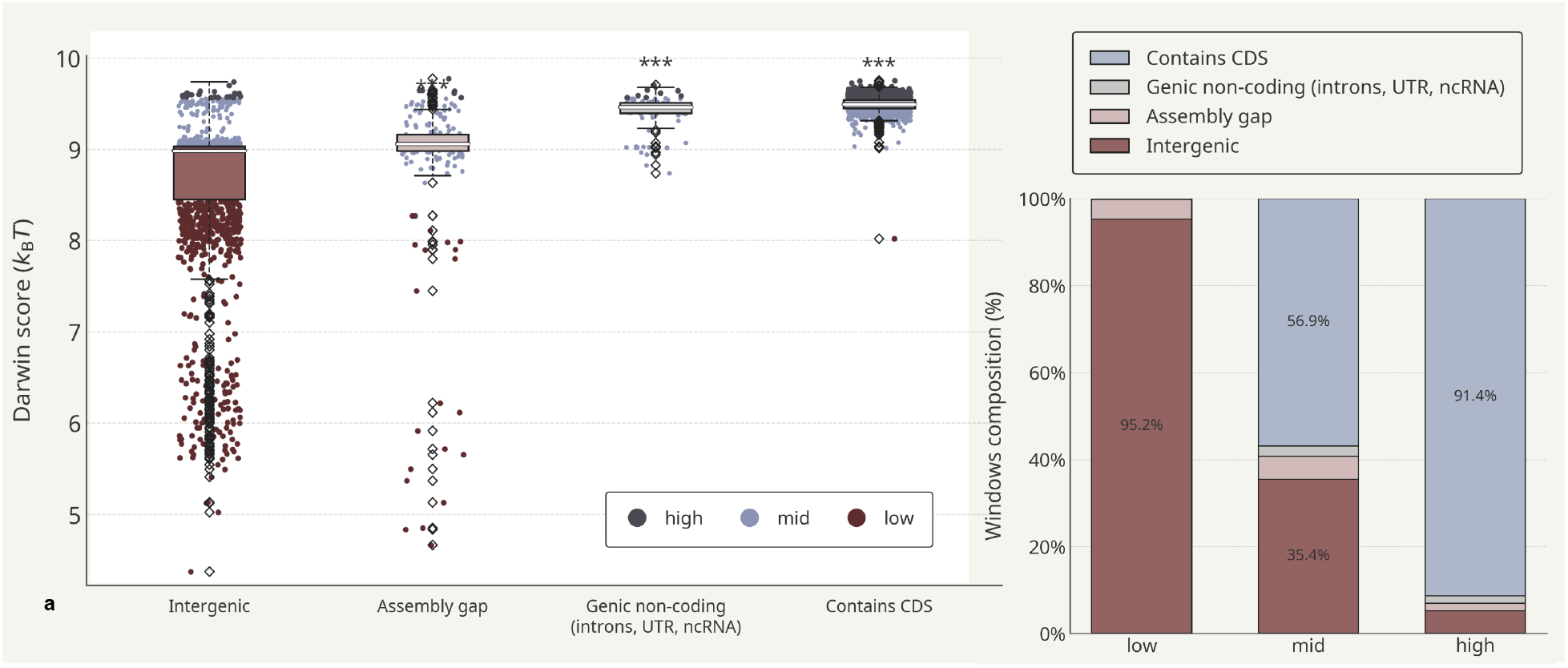
Genome-wide effective-energy deviations in *Drosophila melanogaster*. **a** Distribution of effective-energy deviation scores in non-overlapping 50-kb windows grouped by annotation-overlap class: intergenic, assembly gap, genic non-coding, and contains CDS. Windows were assigned to mutually exclusive classes using the hierarchy Contains CDS *>* Genic non-coding *>* Assembly gap *>* Intergenic. Asterisks indicate significantly higher scores relative to intergenic windows by one-sided Mann–Whitney tests. **b** Composition of annotation-overlap classes within each score category.

The codon analyses use species-wide frequency profiles. As a complementary test of whether sequence composition localizes functional sequences within a genome, we analyzed *D. melanogaster* using a local effective-energy deviation score, defined here as the *Darwin score*. For each chromosome *c*, we estimated the frequency *f*_*c*_(*w*) of each 7-mer and assigned it a dimensionless effective energy

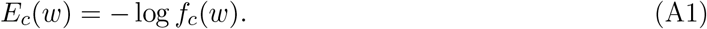

For a genomic region *r*, the score was the average effective energy of its constituent 7-mers,

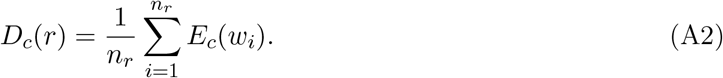

High-scoring windows are enriched in words that are rare relative to the chromosome-wide sequence background. Under GCT, the background distribution is approximately reverse-complement symmetric 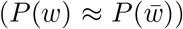; therefore, regions that depart from this background are enriched in words whose frequencies deviate from the RC-balanced expectation. Note that the Darwin score does not directly measure RC asymmetry within a window: two RC partners could both be rare and equally frequent, yielding a high score with intact local symmetry. The score instead captures departure from a chromosomal baseline that GCT predicts to be RC-symmetric, making it an indirect proxy for symmetry breaking at the regional level. This distinction makes the analysis complementary to, rather than redundant with, the codon-level reverse-complement correlations reported in the main text.

The *Drosophila* genome was partitioned into 4,628 non-overlapping 50-kb windows and each window was assigned to an annotation-overlap class (Figure A1). The effective-energy deviation class was strongly associated with functional content (*χ*^2^ = 1424.81, *p* = 2.47 × 10^*−*286^). High-scoring windows were enriched in coding and genic features: 91.4% overlapped coding sequences, 93.1% contained gene annotations, 91.6% included messenger RNA, and 58.3% contained non-coding RNA. Low-scoring windows showed minimal overlap with these features and were dominated by intergenic or weakly annotated regions. Thus, windows enriched in chromosome-rare 7-mers preferentially occur in regions with functional annotation.

## B Mathematical interpretation of the reverse-complement statistic

This appendix gives a compact mathematical reading of the codon-level statistic used in the main text. Its purpose is not to introduce a phenomenological model of codon usage, but only to show that the Pearson correlation used in the exhaustive factorized-involution search has an exact interpretation as a normalized measure of symmetry breaking. The argument has two steps. First, we treat an arbitrary codon-level involution *S*, which includes the factorized transformations used in the exhaustive search. Second, we specialize the same identity to the winning transformation, the reverse-complement map *R*, and write the corresponding correlation as 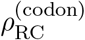.

### B.1 Even–odd decomposition under a codon involution

Let *C* be the set of 64 codons, and let *S* : C → C be any codon-level involution, so that *S*^2^ = id. For a species, let *f ∈* ℝ^64^ be the normalized codon-frequency vector, 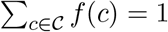 *f* (*c*) = 1, and let *P*_*S*_ be the permutation matrix induced by *S*, (*P*_*S*_*f*)(*c*) = *f* (*S*(*c*)). The empirical statistic used in the main text is

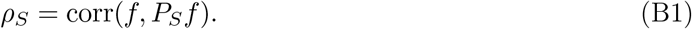

Because *P*_*S*_ is a permutation, *f* and *P*_*S*_*f* have the same mean and variance. Let *µ* = 1*/*64 and define the centered vector *u* = *f* − *µ***1**. Then

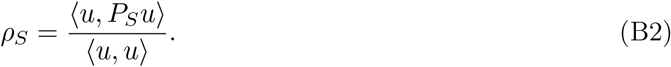

The vector *u* can be decomposed uniquely into components that are even and odd under *S*:

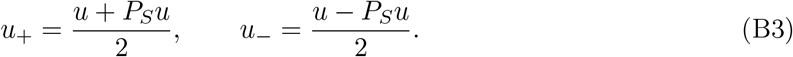

Since *P*_*S*_*u*_+_ = *u*_+_ and *P*_*S*_*u*_*−*_ = −*u*_*−*_, and since the two components are orthogonal,

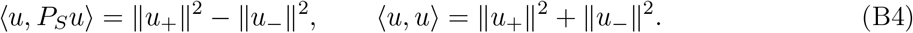

Therefore

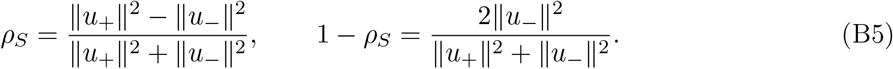

Thus, for every candidate factorized codon involution in the exhaustive search, maximizing *ρ*_*S*_ is exactly equivalent to minimizing the squared component of codon usage that is odd under *S*, normalized by the total centered codon-usage variance. In this sense, the winning transformation is the codon-level symmetry under which the empirical codon-usage vector is least broken. The fact that the winning transformation is the reverse complement means that the least-broken codon-level involution is the one associated with generalized Chargaff symmetry.

Equation B5 also clarifies how to interpret negative correlations. The quantity ∥*u*_*−*_∥^2^*/*(∥*u*_+_∥^2^+ ∥*u*_*−*_∥^2^) is the fraction of the non-uniform codon-usage variance that lies in the *S*-odd subspace. Hence 1 − *ρ*_*S*_ is twice this fraction and lies between 0 and 2. A value *ρ*_*S*_ = −1 means exact symmetry under *S*; *ρ*_*S*_ = 0 means that the even and odd components have equal squared norm; and *ρ*_*S*_ *<* 0 means that the odd component is larger than the even component. In the limiting case *ρ*_*S*_ = −1, all non-uniform variation is odd under *S*.

### B.2 Specialization to the reverse-complement map

We now apply the general identity to the transformation selected by the exhaustive search, namely the reverse-complement map *R*. In this subsection *ρ*_*R*_ is the codon-level reverse-complement correlation used throughout the main text, denoted 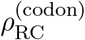.

For *R*, no DNA codon is self-reverse-complementary. A self-reverse-complementary word of odd length would require its central nucleotide to equal its Watson–Crick complement, which is impossible for *A, C, G, T*. Therefore the 64 codons partition into 32 unordered reverse-complement pairs

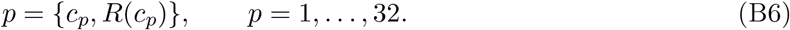

For each pair define the pair weight and pair asymmetry

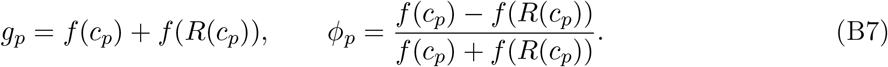

The vector *ϕ* = (*ϕ*_1_, …, *ϕ*_32_) is a codon-pair order parameter for reverse-complement symmetry. Reverse-complement balance corresponds to *ϕ*_*p*_ = 0 for all *p*.

For *S* = *R*, Eq. B5 can be written explicitly in terms of these 32 pair variables. Since the two members of each pair have difference *g*_*p*_*ϕ*_*p*_,

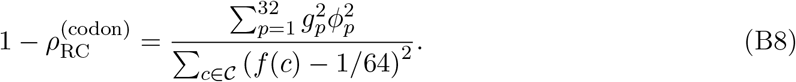

This identity is exact whenever the denominator is nonzero. Consequently, 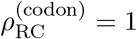 if and only if every reverse-complement codon pair is balanced, *f* (*c*_*p*_) = *f* (*R*(*c*_*p*_)), apart from the trivial uniform profile where the Pearson correlation is undefined. Negative values of 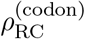 are not exceptional mathematically: they correspond to cases in which more than half of the non-uniform codon-usage variance is reverse-complement-odd. Such profiles are not merely far from reverse-complement balance; they are anticorrelated with their reverse-complement-transformed copies.

Equations B5 and B8 justify the terminology used in the main text. The symmetry being tested is invariance under *R*, and the part that breaks this symmetry is exactly the *R*-odd component. The numerator of Eq. B8 is the total squared reverse-complement codon-pair imbalance, while the denominator is the total centered codon-usage variance. Therefore 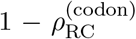 is an exact, scale-free, normalized measure of reverse-complement symmetry breaking: it is zero at perfect reverse-complement balance, increases as pairwise RC imbalance grows, equals one when the RC-even and RC-odd components contribute equally, and exceeds one precisely when the RC-odd component dominates. The full vector *ϕ* retains the pair-specific information that the scalar correlation compresses.

### B.3 Genus-level aggregation and one-species-per-genus bootstrap

Let *G* denote the set of genera and let *g* ∈ *G* contain species *k* ∈ *g*. For each factorized codon involution *S*, the species-level correlation is

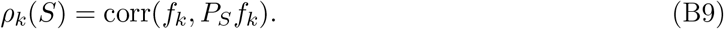

The genus-median version of the transformation score is

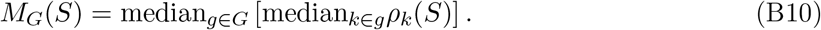

This gives each genus one contribution to the final ranking, rather than allowing species-rich genera to dominate the statistic. Using this genus-level score, the reverse-complement transformation remained the top-ranked factorized codon involution, with *M*_*G*_(*R*) = 0.5220, compared with 0.4854 for the best non-reverse-complement transformation. In an additional bootstrap analysis, one species was sampled at random from each genus and transformations were ranked by the median correlation across the sampled species. The reverse-complement transformation ranked first in all 1,000 bootstrap replicates, with median margin 0.0373 over the best non-reverse-complement transformation. These analyses reduce the possibility that the global reverse-complement optimum is a consequence of uneven taxonomic sampling.

### B.4 Fixed codons and amino-acid preservation among candidate factorized codon involutions

The 39 non-identity factorized codon involutions are not equally favored by trivial self-matching. For a codon transformation *S*, a fixed codon is a codon *c* ∈ C such that

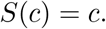

Fixed codons contribute self-comparisons to the Pearson correlation between the codon-frequency vector *f* (*c*) and its transformed version *f* (*S*(*c*)). Thus, transformations with many fixed codons can receive a partial correlation advantage independently of any nontrivial symmetry of codon usage.

We therefore counted, for each candidate factorized codon involution, the number of fixed codons, the number of sense codons mapped to codons encoding the same residue, and the number of codons mapped to the same amino-acid class when stop codons are treated as an additional class. The reverse-complement transformation has zero fixed codons and maps no sense codon to a synonymous codon. By contrast, several competing factorized codon involutions have 8 or 16 fixed codons, and some preserve amino-acid identity for many codons. Therefore, the top rank of the reverse-complement transformation is not explained by trivial codon identity or synonymous preservation.

### B.5 Phylum-level codon reverse-complement symmetry across the tree of life

**Figure S1:**
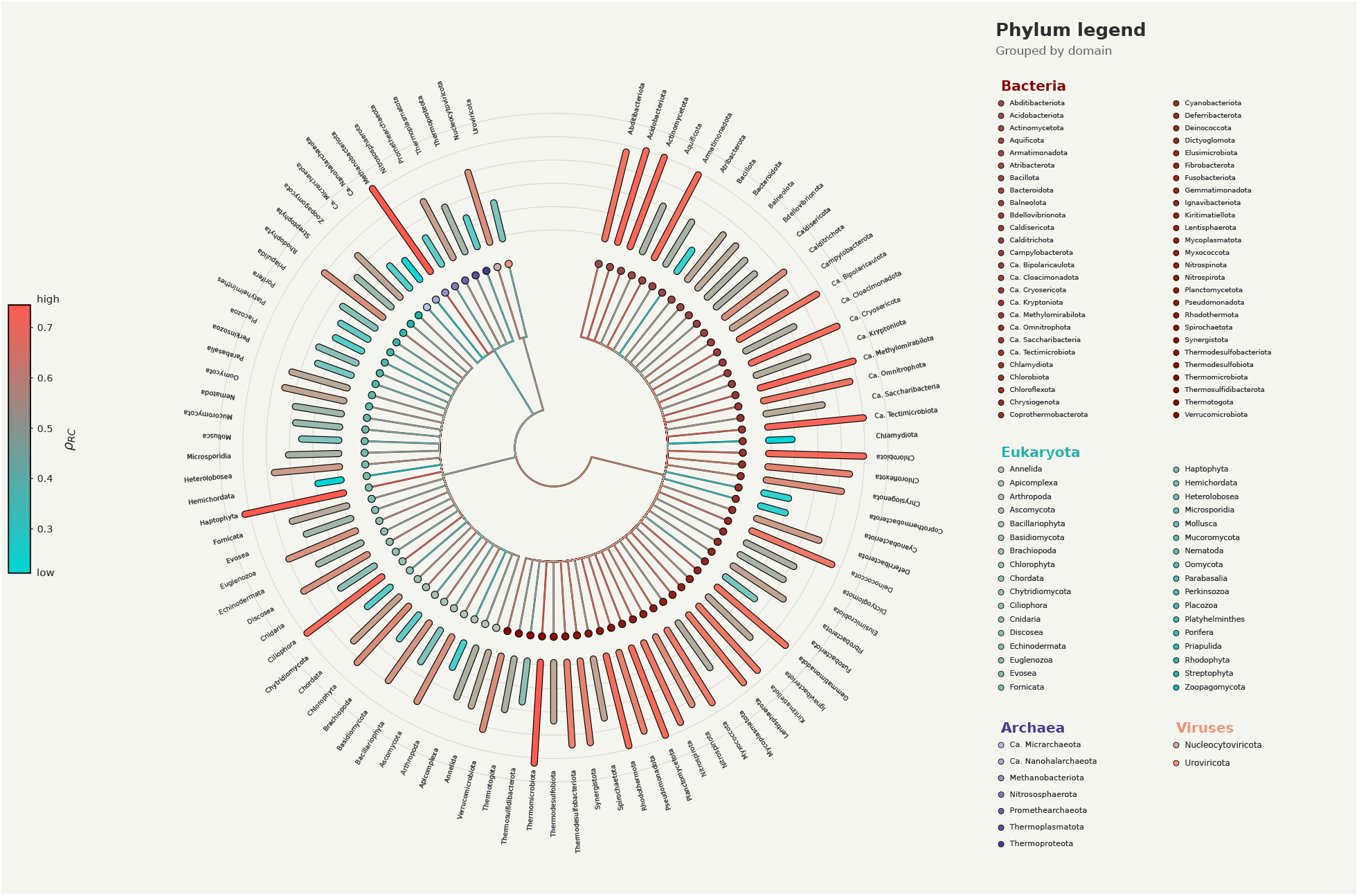
Phylum-level codon reverse-complement symmetry across the tree of life. Radial cladogram of the phyla included in the dataset, grouped by domain (Bacteria, Eukaryota, Archaea, and Viruses; see legend). Each tip is colored according to the median codon-level reverse-complement correlation 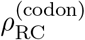 of the species in that phylum (low = teal, high = red). Although all organisms show a positive reverse-complement correlation, *ρ*_*RC*_ *>* 0, its magnitude varies substantially across phyla. This figure summarizes, at phylum-level resolution, the signal quantified at the domain, class, and order levels in Fig. 5. An interactive version is available at https://compbiomed-unito.github.io/chargaff-symmetry-tradeoffs/.

## Data, materials, and software availability

All data used in this study are derived from public codon-usage, genome-sequence, and genome-annotation resources described above. All processed data, taxonomic labels, genome-window scores, randomization scripts, and figure-generation code are available at https://github.com/compbio-med-unito/chargaff-symmetry-tradeoffs

## Author contributions

P.F., L.P., and C.T. conceived the study. P.F., C.T., I.C., V.I., and G.B. performed the computational analyses. All authors contributed to interpretation of the results. V.I., I.C., G.B., C.R., and F.C. assisted with data curation and validation. All authors contributed to writing and critical revision of the manuscript and approved the final version.

## References

[1] C. Alff-Steinberger. “Evidence for a coding pattern on the non-coding strand of the E. coli genome”. In: Nucleic Acids Research 12 (1984), pp. 2235–2241.

[2] D. R. Forsdyke. “Sense in antisense?” In: Journal of Molecular Evolution 41 (1995), pp. 582–586.

[3] D. R. Forsdyke. “Relative roles of primary sequence and (G+C)% in determining the hierarchy of frequencies of complementary trinucleotide pairs in DNAs of different species”. In: Journal of Molecular Evolution 41 (1995), pp. 573–581.

[4] R. Grantham, C. Gautier, M. Gouy, R. Mercier, and A. Pavé. “Codon catalog usage and the genome hypothesis”. en. In: Nucl Acids Res 8.1 (1980), pp. 197–197. DOI: 10.1093/nar/8.1.197-c. URL: https://academic.oup.com/nar/article-lookup/doi/10.1093/nar/8.1.197-c (visited on 02/06/2025).

[5] M. Gouy and C. Gautier. “Codon usage in bacteria: correlation with gene expressivity”. In: Nucleic Acids Research 10.22 (Nov. 1982), pp. 7055–7074. DOI: 10.1093/nar/10.22.7055. URL: https://doi.org/10.1093/nar/10.22.7055 (visited on 12/16/2024).

[6] S. G. E. Andersson and P. M. Sharp. “Codon usage in the Mycobacterium tuberculosis complex”. In: Microbiology 142.4 (1996). Publisher: Microbiology Society, pp. 915–925. DOI: 10.1099/00221287-142-4-915. URL: https://www.microbiologyresearch.org/content/journal/micro/10.1099/00221287-142-4-915 (visited on 12/16/2024).

[7] E. P. Rocha. “Codon usage bias from tRNA’s point of view: Redundancy, specialization, and efficient decoding for translation optimization”. In: Genome Res 14.11 (Nov. 2004), pp. 2279–2286. DOI: 10.1101/gr.2896904. URL: https://www.ncbi.nlm.nih.gov/pmc/articles/PMC525687/ (visited on 12/16/2024).

[8] A. Iriarte, G. Lamolle, and H. Musto. “Codon Usage Bias: An Endless Tale”. eng. In: J Mol Evol 89.9-10 (Dec. 2021), pp. 589–593. DOI: 10.1007/s00239-021-10027-z.

[9] S. T. Parvathy, V. Udayasuriyan, and V. Bhadana. “Codon usage bias”. In: Molecular Biology Reports 49.1 (Nov. 2021), pp. 539–565. DOI: 10.1007/s11033-021-06749-4. URL: http://dx.doi.org/10.1007/s11033-021-06749-4.

[10] Y. Liu. “A code within the genetic code: codon usage regulates co-translational protein folding”. In: Cell Communication and Signaling 18.1 (Sept. 2020), p. 145. DOI: 10.1186/s12964-020-00642-6. URL: https://doi.org/10.1186/s12964-020-00642-6 (visited on 12/16/2024).

[11] P. Fariselli, C. Taccioli, L. Pagani, and A. Maritan. “DNA sequence symmetries from randomness: the origin of the Chargaff’s second parity rule”. In: Brief Bioinform 22.2 (Apr. 2020), pp. 2172–2181. DOI: 10.1093/bib/bbaa041. URL: https://www.ncbi.nlm.nih.gov/pmc/articles/PMC7986665/ (visited on 12/16/2024).

[12] P. Fariselli and A. Maritan. “Thermodynamic perspectives into DNA stability and information encoding in the human genome”. In: Communications Physics 8.1 (Mar. 2025). DOI: 10.1038/s42005-025-02025-0. URL: http://dx.doi.org/10.1038/s42005-025-02025-0.

[13] L. Duret and N. Galtier. “Biased Gene Conversion and the Evolution of Mammalian Genomic Landscapes”. In: Annual Review of Genomics and Human Genetics 10 (2009), pp. 285–311. DOI: 10.1146/annurev-genom-082908-150001. URL: https://doi.org/10.1146/annurev-genom-082908-150001.

[14] F. Lassalle, S. Périan, T. Bataillon, X. Nesme, L. Duret, and V. Daubin. “GC-Content Evolution in Bacterial Genomes: The Biased Gene Conversion Hypothesis Expands”. In: PLOS Genetics 11 (2015), e1004941. DOI: 10.1371/journal.pgen.1004941. URL: https://doi.org/10.1371/journal.pgen.1004941.

[15] C. W. Wheat, H. Vogel, U. Wittstock, M. F. Braby, D. Underwood, and T. Mitchell-Olds. “The genetic basis of a plant–insect coevolutionary key innovation”. In: Proceedings of the National Academy of Sciences 104.51 (Dec. 2007), pp. 20427–20431. DOI: 10.1073/pnas.0706229104. URL: http://dx.doi.org/10.1073/pnas.0706229104.

[16] J. Athey, A. Alexaki, E. Osipova, A. Rostovtsev, L. V. Santana-Quintero, U. Katneni, V. Simonyan, and C. Kimchi-Sarfaty. “A new and updated resource for codon usage tables”. In: BMC Bioinformatics 18.1 (Sept. 2017), p. 391. DOI: 10.1186/s12859-017-1793-7. URL: https://doi.org/10.1186/s12859-017-1793-7 (visited on 12/16/2024).

[17] N. A. O’Leary, M. W. Wright, J. R. Brister, S. Ciufo, D. Haddad, R. McVeigh, B. Rajput, B. Robbertse, B. Smith-White, D. Ako-Adjei, et al. “Reference sequence (RefSeq) database at NCBI: current status, taxonomic expansion, and functional annotation”. In: Nucleic Acids Research 44.D1 (Nov. 2015), pp. D733–D745. DOI: 10.1093/nar/gkv1189. URL: http://dx.doi.org/10.1093/nar/gkv1189.

[18] N. A. O’Leary, E. Cox, J. B. Holmes, W. R. Anderson, R. Falk, V. Hem, M. T. Tsuchiya, G. D. Schuler, X. Zhang, J. Torcivia, et al. “Exploring and retrieving sequence and metadata for species across the tree of life with NCBI Datasets”. In: Scientific data 11.1 (2024), p. 732.

